# Hyperinflammation evokes different antiviral TMPRSS2 and ADAM17 expression responses in human gut xenograft versus host mouse gut although overall genomic responses are similar

**DOI:** 10.1101/2021.05.09.443289

**Authors:** Lorna Morris, Einat Nisim-Eliraz, Iftach Schouten, François Bergey, Nigel Dyer, Hiroshi Momiji, Eilam Nir, Noga Marsiano, Raheleh Sheibani Tezerji, Simcha Yagel, Philip Rosenstiel, David Rand, Vitor A.P. Martins dos Santos, Nahum Y. Shpigel, SysmedIBD research consortium

## Abstract

The global spread of the newly emerged severe acute respiratory syndrome coronavirus 2 (SARS-CoV-2) has led to the pandemic outbreak of coronavirus disease 2019 (COVID-19), an inflammatory disease that is primarily affecting the respiratory system. However, gastrointestinal symptoms in COVID-19 patients suggests that the gut may present another viral target organ. Disease development and severity is dependent on viral interaction with two cell surface human proteins, ACE2 and TMPRSS2, and on antiviral response which may lead to systemic hyperinflammatory syndrome and multiorgan dysfunction. Understanding the host response to SARS-CoV-2 infection and the pathology of the disease will be greatly enhanced by the development of appropriate animal models. Laboratory mice have been the mainstay of therapeutic and vaccine development, however, the virus does not grow in wild type mice and only induced mild disease in transgenic animals expressing human ACE2. As there are known differences between immune response in laboratory mice and humans we evaluated the response of human gut developed as xenografts and host mouse gut following systemic LPS injections as a hyperinflammation model system. The orthologous gene expression levels in the mouse and human gut were highly correlated (Spearman’s rank correlation coefficient: 0.28–0.76) and gene set enrichment analysis of significantly upregulated human and mouse genes revealed that a number of inflammatory and immune response pathways are commonly regulated in the two species. However, species differences were also observed, most importantly, in the inflamed human gut but not in the mouse gut, there was clear upregulation of mRNAs coding for TMPRSS2, ADAM17 and for RIG-I-like receptors, which are involved in the recognition of viruses and in antiviral innate immune response. Moreover, using species-specific immunofluorescence microscopy, we demonstrated the expression and localization of human ACE2 and TMPRSS2 proteins, which are essential elements of the molecular machinery that enables SARS-CoV-2 to infect and replicate in human gut cells. Our findings demonstrate that the intestinal immune response to inflammation in humans and mice are generally very similar. However, certain human-specific diseases, such as COVID-19, can only be successfully studied in an experimental model of human tissue, such as the gut xenograft.

## INTRODUCTION

The global spread of the newly emerged severe acute respiratory syndrome coronavirus 2 (SARS-CoV-2) led to pandemic outbreak of human coronavirus disease 2019 (COVID-19), an influenza-like disease that is primarily affecting the respiratory system. However, COVID-19 patients often present with gastrointestinal symptoms such as diarrhea, vomiting, and abdominal pain. suggesting that the intestine may present another viral target organ (1). Two cell surface proteins; angiotensin-converting enzyme 2 (ACE2) and transmembrane serine protease 2 (TMPRSS2) constitute the molecular machinery enabling SARS-CoV-2 to infect and replicate in human cells. However, recent studies suggest that additional host factors such as the membrane protein neuropilin-1 (NRP1) and the proteases furin and metalloproteinase domain 17 (ADAM17) might also be involved in SARS-Cov-2 processing and entry in human cells (2–6).

Severe COVID-19 disease is often associated with systemic hyperinflammatory syndrome and multiorgan dysfunction reminiscent of sepsis (7). Understanding the host response to SARS-CoV-2 infection and the pathology of disease will be greatly enhanced by the development of appropriate animal models. Laboratory mice have been the mainstay of therapeutic and vaccine development, however, the virus does not grow in wild type mice and only induced mild disease in transgenic animals expressing human ACE2 (8). Furthermore, the validity of mice as a model system in translational research was seriously questioned by Seok et al reporting that transcriptional responses in mouse models poorly mimic human inflammatory diseases (9). However, alternative analysis of the same gene expression data sets was suggested by Takao and Miyakawa (10) and led to a diametrically opposite conclusion that genomic responses in mouse models greatly mimic human inflammatory diseases. For obvious reasons, this study and many current COVID-19 studies were based on comparative transcriptome analysis of blood leukocytes derived from human patients affected by inflammatory conditions compared with the corresponding mouse models. The use of blood leukocytes as surrogates for transcriptome analysis of inaccessible human tissues in patients was criticized by several authors suggesting the use of parenchymal cells of affected organs for comparative transcriptome analysis in inflammatory conditions (11, 12). Nevertheless, most of our understanding of disease mechanisms and biology in inflamed organs is based on systematic observations in experimental animals and episodic studies of humans; there is a dearth of comparative transcriptomic data about the human and mouse gut in hyperinflammatory conditions. Here we used human gut tissues developed in mice as xenografts to study the expression and localization of SARS-CoV-2 entry mechanisms and whole genome gene expression response in the human gut under hyperinflammatory conditions.

We have utilized an experimental platform, first reported by Winter et al in 1991 (13) and refined in our laboratories for study of enteric nervous system (14, 15), human-specific pathogens (16–18) and IBD (19–22). Fetal gut is obtained from pregnancy terminations performed legally at 12-18 weeks gestational age and transplanted subcutaneously in mature SCID mice, where it grows and can be experimentally manipulated over the course of the subsequent several months. Transplanted mice were subjected to systemic LPS, a well-established model of experimental systemic hyperinflammatory disease (23–26).

Gene expression profiles were analyzed in human gut transplants and host mouse gut originating from normal control and LPS-treated mice. Next, we adopted the strategy and methodology suggested by Takao and Miyakawa (10) and focused on orthologous genes whose expression levels were significantly changed in both human and mouse gut and used Spearman’s rank correlation for comparison of interspecies agreement. Further, we utilized the ImmuneSigDB database (27), a compendium of approximately 5000 gene expression signatures generated by 389 published studies, to compare the functional consequences of gene expression in mouse and human gut transplants and provide insight into both conserved and species-specific transcriptional responses. Gene Set Enrichment Analysis (GSEA) is used to identify functionally related programs of gene expression (20) for differentially expressed genes using different gene sets, for example Gene Ontology or ImmuneSigDB. GSEA enables relatively small changes in gene expression in a large number of genes to be taken into account when exploring changes in cell state. Genes can be ranked according to a variety of metrics, such as their log fold change in expression or a signal-to-noise calculation, using the mean and variance estimates for the expression of each gene (20). The leading edge genes in a GSEA analysis are the genes whose expression profile drives the GSEA enrichment statistic and thus are those that appear at the top of the ranked list of differentially expressed genes provided to the GSEA algorithm. By systematically comparing the leading edge genes of the enriched gene sets in the mouse and human we defined a gene expression signature consisting of anti-viral response genes that was specific for hyperinflammation in the human gut.

## RESULTS

### ACE2 and TMPRSS2 are expressed in the human gut xenograft

Using immunofluorescence microscopy, we first analyzed human fetal gut (Fig. S1) and mature human gut xenografts for the expression and cellular localization of ACE2 and TMPRSS2 (Fig. 1). Here we show that in steady state human gut xenografts ACE2 and TMPRSS2 are exclusively expressed by gut epithelial cells. However, while all gut epithelial cells were highly positive for cytoplasmic TMPRSS2, ACE2 was specifically localized to the apical membranes along the brush borders of villi epithelial cells. The fact that these cells express both ACE2 and TMPRSS2 indicate susceptibility for SARS-CoV-2 infection and suggest that luminal SARS-CoV-2 viral particles will be able to infect the epithelial cells of the human gut transplants. It is now well established that SARS-CoV-2 infection of the gut does not recapitulate the severe disease and pathology inflicted by respiratory infection which is frequently associated by systemic hyperinflammatory immune response. To better understand and compare human and mouse gut response to hyperinflammation, we next used whole-genome transcription analysis of human gut xenografts and host mouse gut hyperinflammatory response to systemic LPS.

**Fig. 1.**
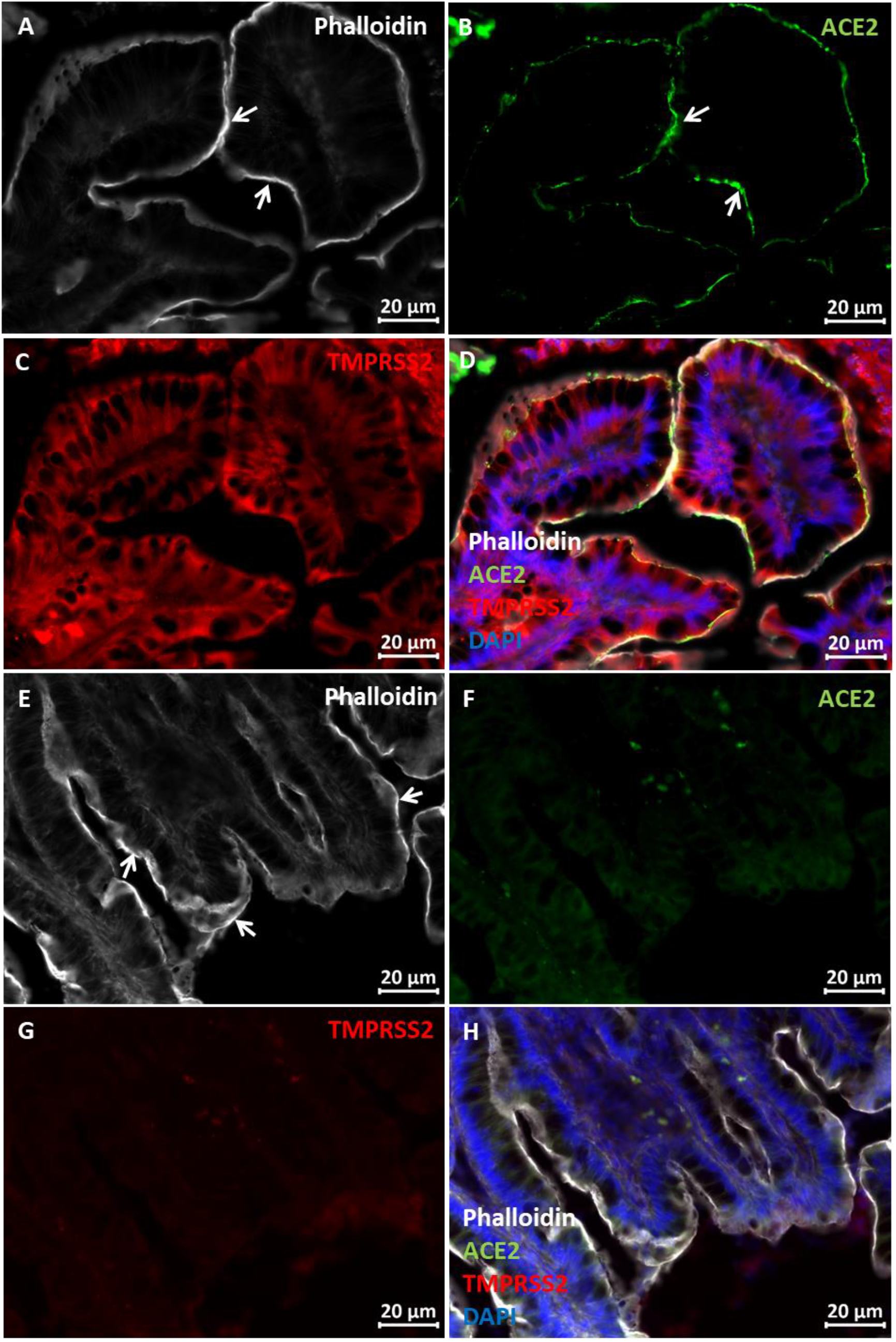
SARS-CoV-2 entry molecule; ACE2 and TMPRSS2 are co-expressed in the mucosal villi of human gut xenografts. Fluorescence microscopy was used to analyze cryosections of normal human gut xenografts stained using DAPI, phalloidin and anti ACE2 and TMPRSS2 antibodies (A-D). Images of no-primary controls are shown in E-H. Microvilli of gut epithelial cells are demon-strated by phalloidin staining (white arrows in A&E) co-localized with ACE2 staining (white arrows in B). Scale bars 20 μm.

### High Spearman Correlation of differential gene expression in hyperinflammation between Human and Mouse Gut

The differential gene expression between normal and inflamed human gut transplants and between normal and inflamed mouse gut are presented in Fig. 2 and Fig. 3, respectively. Differential expression (log_2_ fold-changes) in septic gut of human and mouse orthologs were compared. First, all significantly changed orthologous genes (*P*_adj_-value < 0.05; |log_2_ fold-change| >1) in either human or mouse gut were compared using Pearson’s correlation coefficient R^2^ (Fig. 4). Non-significant log_2_ fold-changes were assigned the value of 1 (log2 = 0 in Fig. 4) and are located on the y-axis (for non-significant human genes in Fig. 4) or on the x-axis (for non-significant mouse genes in Fig. 4). This analysis recapitulated the data analyses performed by Seok et al (9) suggesting poor correlation between the genomic response in septic human and mouse gut (Fig. 4; Pearson’s correlation coefficient R^2^: 0.0025 – 0.0724).

**Fig. 2.**
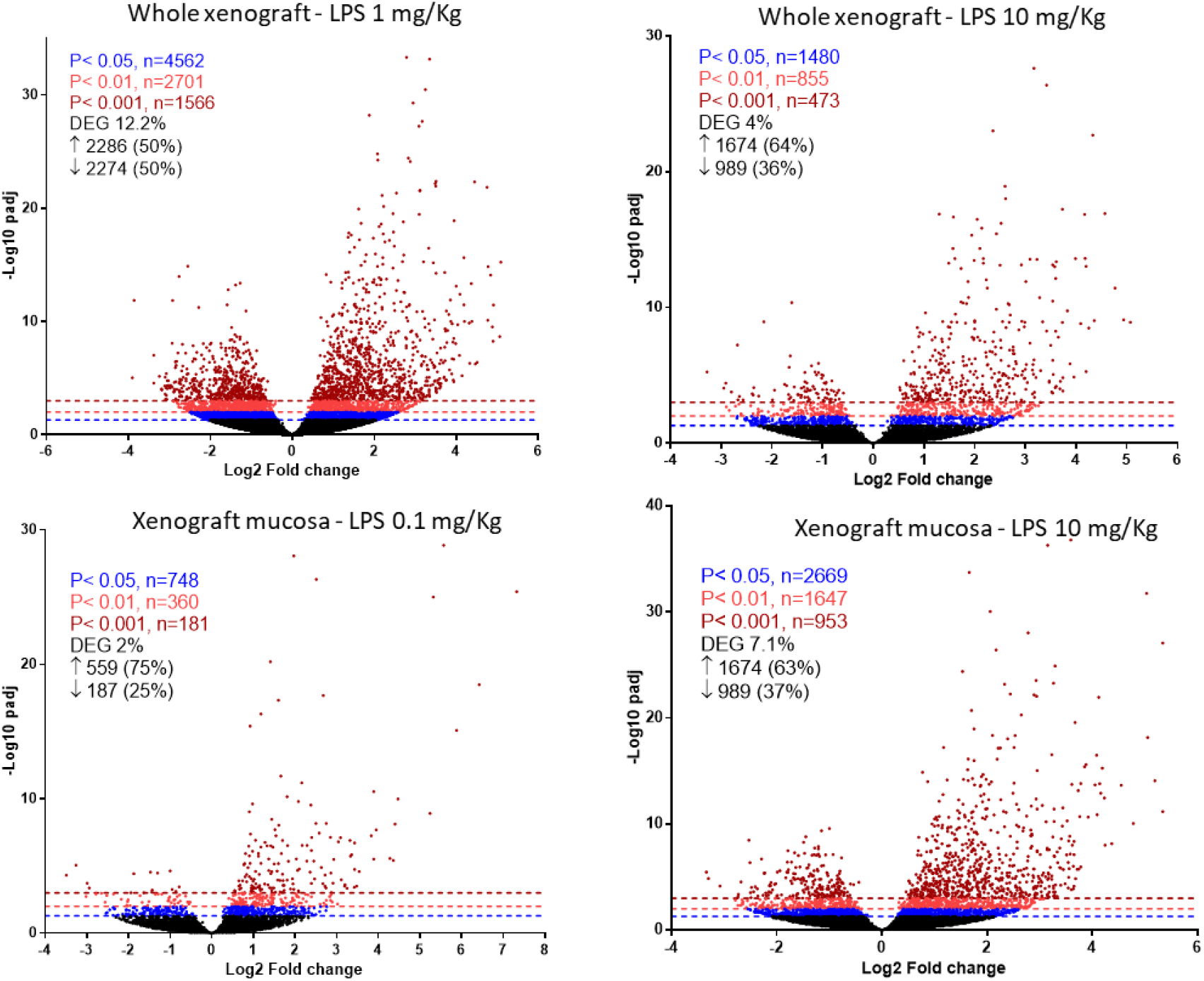
Volcano plots showing fold-change of gene expression in LPS-treated human gut xenografts compared to PBS-treated xenografts. Mice were systemically treated with 0.1, 1 and 10 mg/kg LPS and total RNA was extracted from whole xenografts or mucosal scrapings. The x axis indicates the log2 differences in gene-expression level between LPS and PBS treatment groups, with larger positive values representing genes with higher expression in LPS-treated xenografts relative to the PBS-treated xenografts and larger negative values representing genes with higher expression in the PBS-treated xenografts relative to the PBS-treated xenografts. The y axis shows the −log10 of the adjusted P values for each gene, with larger values indicating greater statistical significance. Those genes showing significantly different expression between the two groups are highlighted; P < 0.05 in blue, P < 0.01 in green and P < 0.001 in brown with dashed horizontal lines corresponding to these P values on the y axis. Black points represent genes for which there was no significant difference in gene expression.

**Fig. 3.**
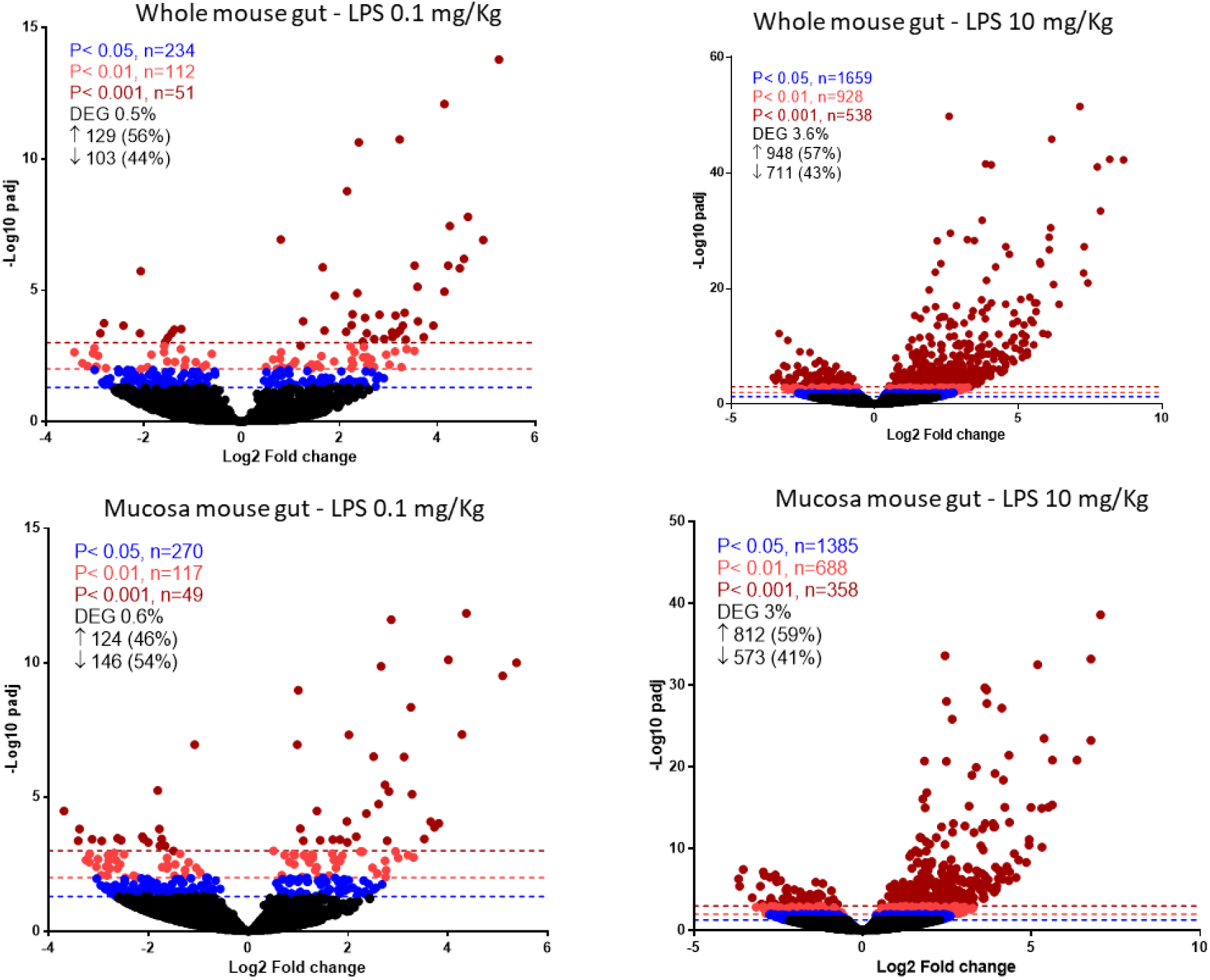
Volcano plots showing fold-change of gene expression in LPS-treated mouse gut compared to PBS-treated mice. Mice were systemically treated with 0.1 or 10 mg/kg LPS and total RNA was extracted from whole xenografts or mucosal scrapings. The x axis indicates the log2 differences in gene-expression level between LPS and PBS treatment groups, with larger positive values representing genes with higher expression in LPS-treated xenografts relative to the PBS-treated xenografts and larger negative values representing genes with higher expression in the PBS-treated xenografts relative to the PBS-treated xenografts. The y axis shows the −log10 of the adjusted P values for each gene, with larger values indicating greater statistical significance. Those genes showing significantly different expression between the two groups are highlighted; P < 0.05 in blue, P < 0.01 in green and P < 0.001 in brown with dashed horizontal lines corresponding to these P values on the y axis. Black points represent genes for which there was no significant difference in gene expression.

**Fig. 4.**
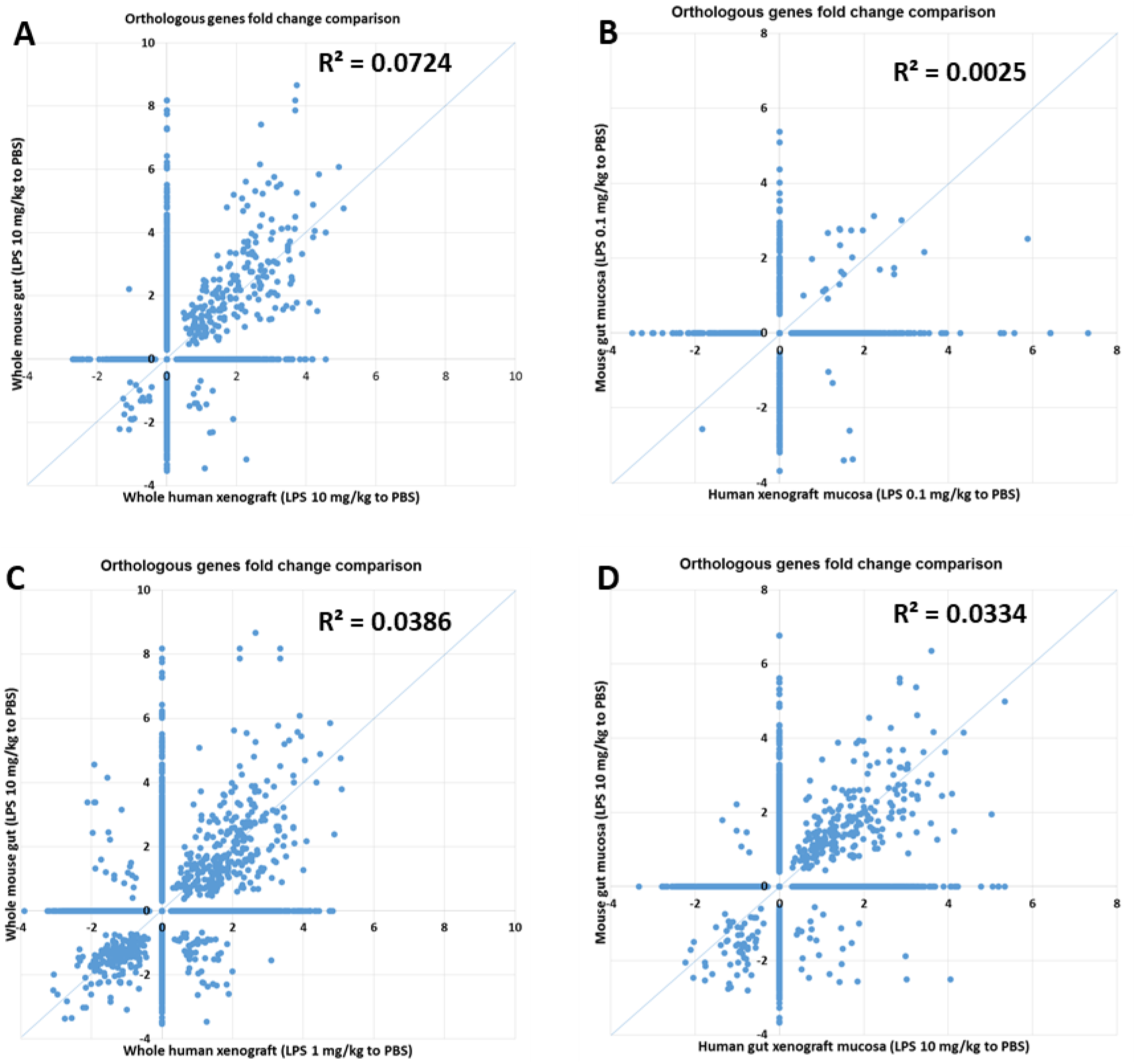
Differentially expressed genes (DEG) shown in Fig. 1 and Fig. 2 were used to correlate the magnitude of gene expression in response to LPS between the mouse (y axis) and human gut (x axis). Vertical bar and horizontal bar for each scatter plot represents log2 fold change (FC) in mouse and human gut, respectively. Genes with log2FC values >0 were more highly expressed in LPS-treated gut relative to PBS-treated gut, and genes with log2FC values < 0 were more highly expressed in PBS-treated gut relative to LPS-treated gut. Scatter plots and Pearson’s correlation coefficient R^2^ of significantly expressed gene changes (P < 0.05; |FC| >1) in response to LPS treatment in either human or mouse gut. Non-significant FC were assigned the value of 1 (log2 = 0) and are located on the y-axis (for non-significant human genes) or on the x-axis (for non-significant mouse genes).

We compared genes whose expression levels were significantly changed in both human and mouse gut (*P*_adj_-value < 0.05; |log_2_ fold-change| >1) using Spearman’s rank correlation coefficient ρ (Fig. 5). Genomic response of significantly expressed orthologous genes in the septic human and mouse gut was highly correlated (Fig. 5; Spearman’s rank correlation coefficient: 0.328-0.762; P = 0.0388 to P < 0.0001). Consistently high correlations were observed between gene expression in the mouse gut following high dose LPS treatment (10 mg/kg) and gene expression in the human gut at all LPS dosage levels, showing that a high dose of LPS was required to elicit a response in the mouse gut (10mg/kg), whilst even low doses of LPS (0.1mg/kg) elicited a response in the human gut (Fig. S2).

**Fig. 5.**
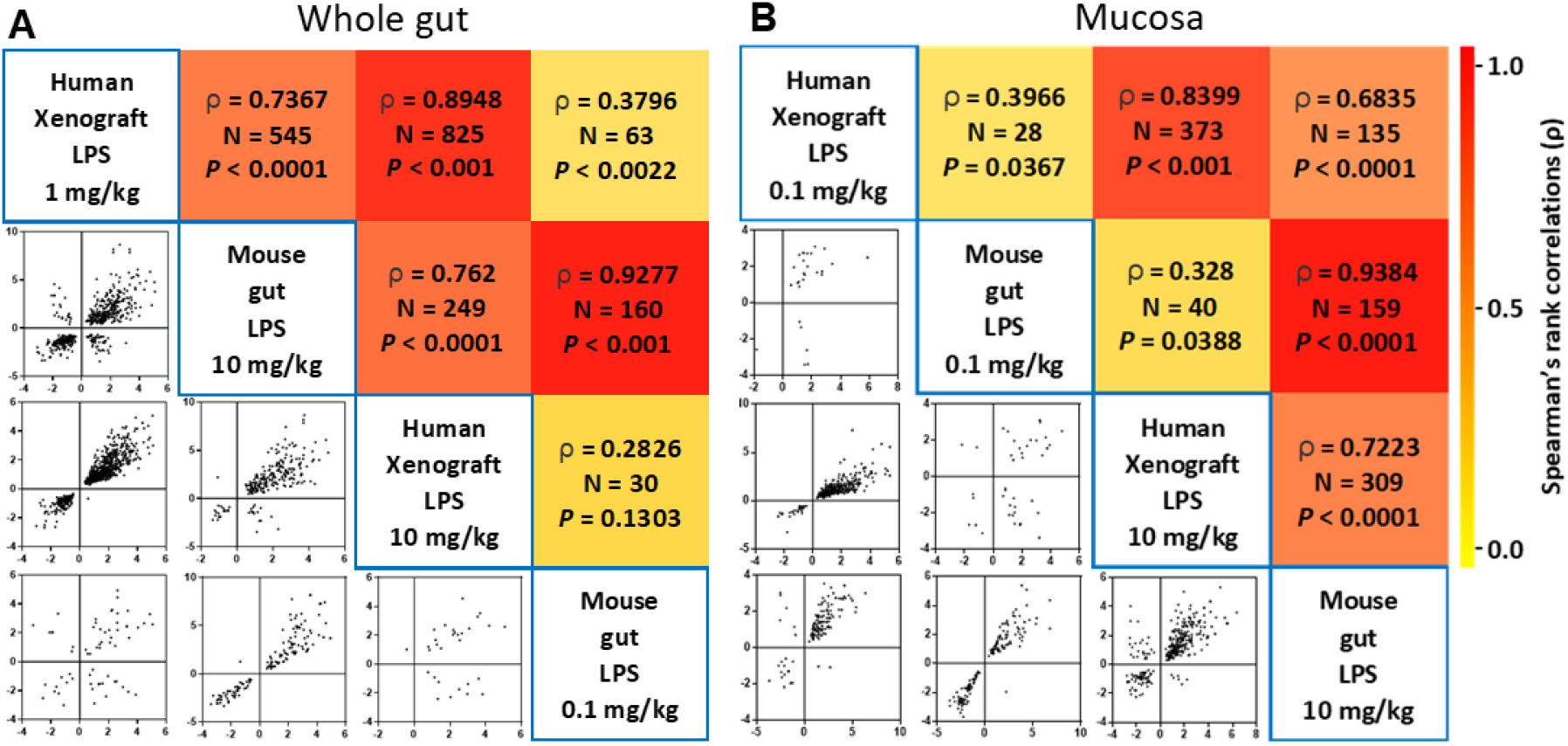
Differentially expressed genes (DEG) shown in Fig. 2 and Fig. 3 were used to correlate the magnitude of gene expression in response to LPS between the mouse (y axis) and human gut (x axis). Vertical bar and horizontal bar for each scatter plot represents log_2_ fold change (FC) in mouse and human gut, respectively. Genes with log2FC values > 0 were more highly expressed in LPS-treated gut relative to PBS-treated gut, and genes with log2FC values < 0 were more highly expressed in PBS-treated gut relative to LPS-treated gut. Scatter plots and non-parametric Spearman rank correlation coefficient ρ of gene changes significantly expressed (P < 0.05; |FC| >1) in response to LPS treatment in both human or mouse gut.

Our approach above followed that of Takao and Miyakawa (10) in using the Spearman correlation coefficient rather than the Pearson coefficient. As pointed out in (10), the Spearman coefficient is a nonparametric approach to identifying a monotone relationship between two sets of data and, unlike the Pearson coefficient, is not based on normality of the data and is generally much more robust. In addition to the better robustness and the non-normality of our data, the Spearman coefficient is also more appropriate to our task which is to determine the strength of any monotone relationship between the two data sets rather than to quantify a linear relationship between them. Thus, given the much greater appropriateness and robustness of the Spearman coefficient our data suggests highly correlated gene expression in response to hyper-inflammation in the mouse and human gut.

### Conserved enrichment of immune-specific Gene Sets in hyper-inflamed Human and Mouse Gut

To probe the functional consequences of the observed gene expression responses to hyper-inflammation and further test the level of correlation we performed GSEA to compare enrichment of functionally related gene sets and the leading edge genes in the mouse and human responses (28). We reasoned that sub-groups of genes that are co-regulated are likely to represent distinct biological processes or pathways and that comparison of these across mouse and human is another method to address comparability of the transcriptional response to hyper-inflammation.

We performed GSEA using the differential expression results at the LPS concentration of 10mg/kg in both human and mouse (compared to the PBS control) as this treatment had the most highly correlated response between orthologs and more differentially expressed genes than at the lower concentrations of LPS (Fig. 5 and Fig. S2). GSEA can be performed with different collections of gene sets, for example Molecular Signatures Database (MSigDB), Gene Ontology and a variety of other databases containing data describing biological pathways (29, 30). Recently Godec et al (27) created the ImmuneSigDB collection, approximately 5,000 gene sets derived from 389 immunological studies. The sets are experimentally derived and include different cell lineages, states and treatments (including LPS stimulation). The authors showed a highly conserved pattern of gene expression in response to sepsis in mouse and human, but also a species-specific response in distinct biological processes. We ranked all significantly differentially expressed orthologous genes (*P*_adj_-value <0.05) according to their fold changes (LPS-stimulated vs unstimulated) in each data set (mouse or human) and performed GSEA using the ImmuneSigDB gene sets. The most enriched gene set in our model system of the human gut gene expression originated from a study in which monocytes were isolated from healthy human donors and were treated with either LPS or vehicle and the gene expression profile compared (GSE9988 with ES= 0.835, adjusted p-value =0.0009). This gene set was also highly enriched when we performed the GSEA with our mouse gut differential expression data (ES= 0.800, adjusted p-value =0.001). This is consistent with the findings of Godec *et al*, showing that the transcriptional response to sepsis is highly conserved.

Fig. 6 shows the GSEA cumulative enrichment scores of the top five enriched gets sets in human (Fig. 6A) compared with the same gene sets in mouse and the top five enriched gets sets in mouse (Fig. 6B) compared with the same gene sets in human. The enrichment profiles and peak scores are very similar and the most highly enriched gene sets are common between mouse and human. The leading edge genes in a GSEA analysis are the genes whose expression profile drives the GSEA enrichment statistic and thus are those that appear at the top of the ranked list of differentially expressed genes provided to the GSEA algorithm. In both mouse and human gut the majority of the most highly enriched gene sets are from experiments in which monocytes, macrophages or dendritic cells were treated with LPS. These genes sets predominantly contain genes coding for pro-inflammatory cytokines including C-X-C motif chemokines, interleukins, cell adhesion proteins and regulators of the adaptive and innate immune response.

**Fig. 6.**
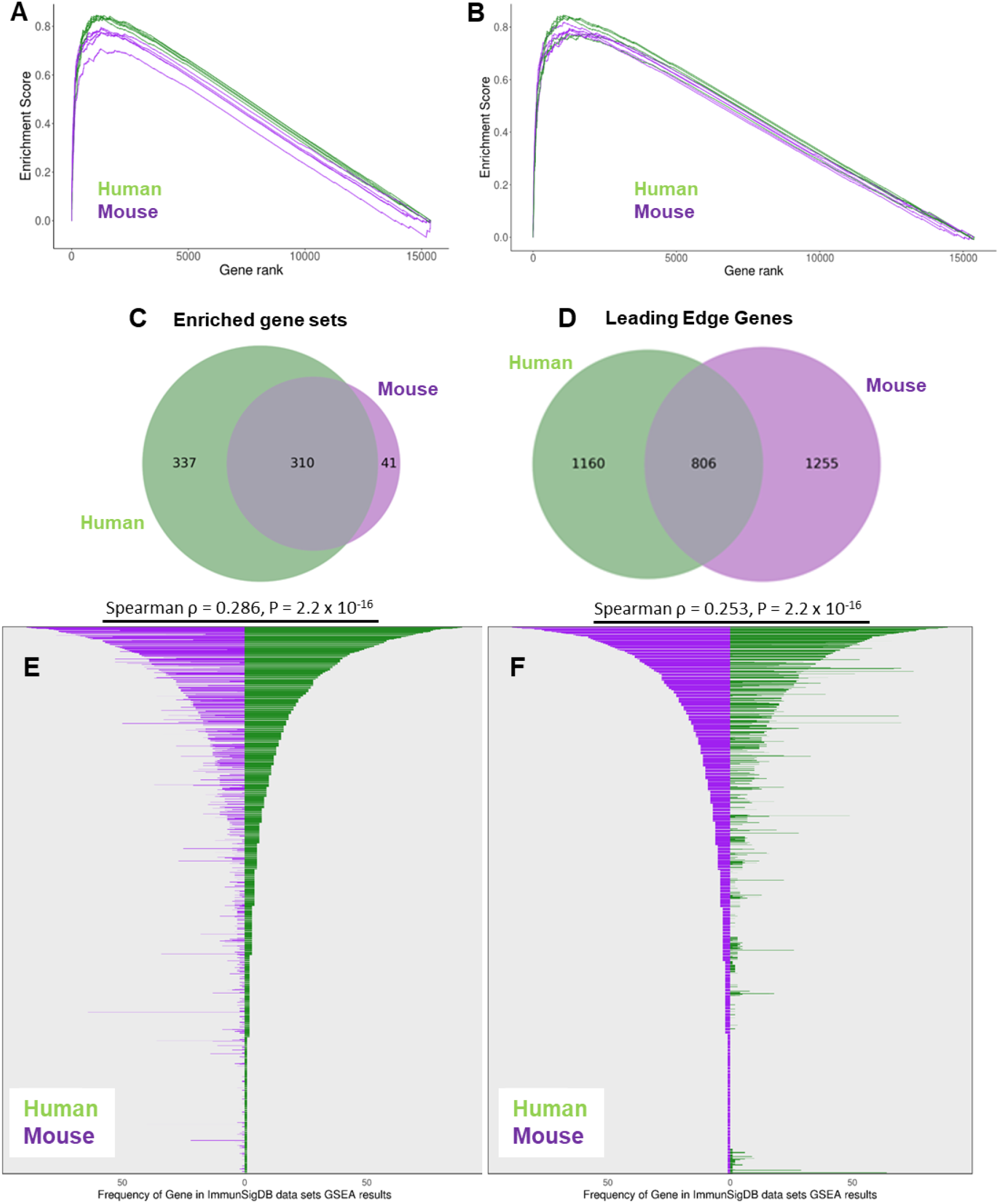
Transcriptional programs are conserved in sepsis between mouse gut and human in gut xenografts. GSEA using ImmunSigDB for genes differentially expressed in LPS treated (10 mg/kg) vs untreated (PBS), whole gut and mucosa combined, with genes ranked by differential expression. For the input to the GSEA the best match orthologs (18,742 genes) are used (using Ensembl Biomart version 83). Mouse results are shown in purple and human in green. (A) The Gene Sets with the 4 highest Enrichment Scores from the GSEA results when ranked by MOUSE Differential expression are displayed (and the GSEA result from the equivalent experiment in human is compared). (B) The Gene Sets with the 4 highest Enrichment Scores from the GSEA results when ranked by HUMAN Differential expression are displayed (and the GSEA result from the equivalent experiment in mouse is compared). (C) Venn diagram to show the overlap in enriched Gene Sets between human and mouse and the overlap in number of genes enriched in those Gene Sets. (D) Venn diagram to show the overlap in number of leading edge genes enriched in the common Gene Sets between mouse and human. Leading edge genes differentially expressed in mouse and human gut are conserved in enriched gene sets from ImmuneSigDB Gene. (E) Comparison of the frequency of genes occurring in the leading edge of the most enriched gene sets from human with mouse. (F) Comparison of the frequency of genes occurring in the leading edge of the most enriched gene sets from mouse with human.

Fig. 6C shows the overlap in enriched Gene Sets between human and mouse, indicating 310 Gene Sets were enriched in both species. Of these 310 enriched Gene Sets, there are 806 leading edge genes common to both species (Fig. 6D). We compared the leading edge genes between the enriched gene sets in our mouse and human comparison and calculated the frequency of occurrence of the genes in these enriched Gene Sets. Fig. 6E-F shows the frequency histograms of leading edge gene occurrences ordered either by their frequency in human (Fig. 6E) or their frequency in mouse (Fig. 6F).

The symmetrical distribution (Fig. 6E-F) in the mouse and human histogram comparison are indicative of the conserved differential expression in these gene signatures, and also supported by the good correlation in log fold changes (ρ = 0.253, p-value = 2.2e-16 for mouse, ρ = 0.286 p-value = 2.2e-16 for human). There is a higher correlation between the most frequently occurring genes in the enriched Gene Sets (Fig S3). Scatterplots to compare the log fold changes in human and mouse of the most frequently occurring leading edge genes are also shown in Figure S3A-B, highlighting those genes that are differentially expressed in mouse but not in human. The human genes with log2 fold changes between 0 and 1 are generally not significant (p adj > 0.05) but several of these genes are more highly expressed in mouse (with log2 fold changes between 1 and 5) and are significant. Fig. S3C-D also shows the most frequently occurring genes, highlighting the similarity and differences between the species at the individual gene level.

### Identification of species-specific components of transcriptional response to hyper-inflammation reveals differences in anti-viral response genes in human xenograft and mouse gut

In general, the leading edge genes of the enriched gene sets are similar for both species, as shown by the overall symmetrical figures (Fig. 6E-F and Fig. S3C-D). However, there are several exceptions where genes are up-regulated in human but not significantly differentially expressed in mouse, and thus do not appear in the leading edge list of the mouse gene sets. These are shown by the shorter or sometimes absent bars for the mouse genes and include the antiviral response genes; IRF7, OAS2, ISG15, DHX58, DDX58, IFIH1, IFI44 (DDX58/IFIH1-mediated induction of interferon-alpha/beta pathway), TRAF1 and CFLAR (apoptosis) EHD1, PTGER4, ETS2, ACSL1, RNF19B, PELI, NAMPT (Fig. S3B and C). These up-regulated human genes are annotated with several GO terms, including GO:0016032 (Viral Process), GO:0045087 (Innate immune response), GO:0051607 (Defense response to Virus) and GO:0032480 (Negative regulation of Type I interferon production).

Similarly, analysis of the mouse leading edge genes revealed up-regulated genes in the septic mouse gut with orthologs that were not significantly differentially expressed in the septic human gut. These include GBP2, CASP4, MARCKSL1, RTP4, IFITM3, CLIC4, MYD88, PROCR, WARS (Fig S2A and D). IFITM3 and the other members of the IFITM (interferon-induced transmembrane family) are known to have anti-viral properties (31). IFITM3, MYD88, CASP4, GBP2 are annotated with GO terms GO:0045087 (Innate immune response) and GO:0060337 (Type I Interferon signaling). Thus human and mouse gut gene expression responses to hyper-inflammation results in distinct innate immune response and Type I interferon signaling.

As we observe differences in expression of the anti-viral response genes between LPS treated human gut and LPS treated mouse gut we reasoned that other important genes in the response to virus may differ between the species that did not appear in the most significant GSEA results. Therefore, we focused our attention on the top 200 up-regulated genes (log2 FC > 1.5 with the lowest adjusted p-values) in each species that were not significantly up-regulated (log2 FC <0.5 and/or p adj >0.05) in the other species and searched their GO Biological Process annotations for virus-related GO terms. In the top 200 human up-regulated genes there are 89 genes that are not significantly differentially expressed in mouse, 12 of which are annotated with GO terms related to response to virus. Of particular interest is the serine protease TMPRSS2 annotated with GO term – GO:0046596 Regulation of Viral Entry into the Host Cell. Recent publications report the importance of TMPRSS2 in spike protein priming of SARS-CoV-2 and show that SARS-CoV-2 infection is enhanced by TMPRSS2 (32, 33). TMPRSS2, ACE2, NRP1, FURIN and ADAM17 have recently been suggested to be involved as SARS-CoV-2 processing and entry in human cells (2–6). Although all these genes are highly expressed in both the mouse and human gut, ACE2, NRP1 and FURIN are not up-regulated in response to LPS in our experiments, whilst TMPRSS2 and ADAM17 are strongly up-regulated in septic human gut (log2 FC 2.73, p adj < 0.000001 and log2 FC 1.06, p adj < 0.00001, respectively) but not in mouse gut (log2 FC 0.5, p adj = 0.39 and log2 FC 0.38, p adj = 0.32, respectively) (Fig. S4).

## DISCUSSION

Along with the respiratory tract, there are increasing data showing that the gastrointestinal tract might also represent a target organ of SARS-CoV-2, and that infected patients could have corresponding organ damage and symptoms (34). We show here that molecular components of the SARS-CoV-2 processing and entry machinery; ACE2, TMPRSS2, NRP1, Furin and ADAM17 (2) are highly expressed in steady-state and hyper-inflamed human gut xenografts. The fact that human gut cells in the transplants express combinations of these molecules indicate susceptibility for SARS-CoV-2 infection and suggest that luminal SARS-CoV-2 viral particles will be able to infect and replicate in the epithelial cells of the human gut transplants (32, 33). Moreover, we speculate that the up-regulation of TMPRSS2 and ADAM17 in response to hyper-inflammatory immune response in the human gut might make these cells more susceptible to SARS-CoV-2 infection, and therefore a better *in vivo* model to study the disease than mouse. Recent studies have shown that local and systemic hyper-inflammation caused by excessive immune responses to infection by SARS-CoV-2 was associated with severe disease and increased mortality (35, 36). Administration of systemic LPS is a well-established hyper-inflammation model. We show here that LPS-induced hyper-inflammation elicits highly similar genomic response in the human and mouse gut. The majority of protein-coding differentially expressed genes, with the highest fold changes in expression in response to LPS have orthologs that are highly correlated between both species. The conserved response to LPS between human and mouse gut was found in 310 gene sets from the ImmuneSigDB collection composed of experimentally derived gene signatures including different cell lineages, states and treatments. However, in these enriched gene sets some differences were observed between the human and mouse gut response to hyper-inflammation, most notable were the increased genetic response of TMPRSS2, ADAM17 and the retinoic-acid-inducible gene I (RIG-I)-like receptors (RLRs) that were not observed in the mouse gut. The later is a family of cytosolic pattern recognition receptors (PRRs) that recognize viral nucleic acid and activate antiviral innate immune response. The RLR comprises the retinoic-acid-inducible gene I (RIG-I)-like receptors (RLRs), encompassing RIG-I (also known as DDX58), melanoma differentiation-associated gene 5 (MDA5; also known as IFIH1) and laboratory of genetics and physiology 2 (LGP2; also known as DHX58), which are ubiquitously expressed in the cytoplasm (37). All these PRRs signaling pathways converge with the activation of the transcription factor NF-kB, the master regulator of inflammation and the transcription factors that control interferon (IFN) induction, IRF3 and IRF7. While recognition of gut-resident viruses by gut PRRs (TLRs and RLRs) contributes to maintaining gut homeostasis (38–41), these same mechanisms are also involved in the pathogenesis of various vital diseases (42). Considering the pivotal role of type I interferon response in the pathogenesis of COVID-19 disease (43–45) and the species differences we show here, human cells and tissues are probably essential for COVID-19 research. The use of human fetal lung transplant in mice (46) and human gut organoids (47) as model systems for SARS-CoV infection were recently reported. Similarly, we suggest that human gut xenografts in mice will be an excellent platform to study the pathogenesis of COVID-19 and for the development of novel therapeutics and vaccines. The human gut tissue vascularizes, expands and persists as a human gut implant, and can be experimentally manipulated over the course of the subsequent several months. While ectopic and not functional, these gut implants develop characteristic structures of the human gut with extensive vasculature and their structural features are highly similar to those of normal human gut, including mucosal villous epithelium, crypt structures, blood vessels and enteric nervous system.

We and others have previously shown that virtually all the cell types that are present in the normal human gut are also present in these xenografts (14, 16–18, 21, 48, 49). Moreover, the general architecture of the gut appears normal, and the tissue is well-vascularized by a human capillary system that anastomoses to the circulatory system of the murine host. In this, the experimental platform is similar to other examples of development of human tissues (lung, skin and liver) subcutaneously transplanted into SCID mice (46, 50). Furthermore, we have shown that many human innate and adaptive components of immune system, which have been shown to be already active in fetal gut at the time of transplantation (51–54), are present and active in the mature xenograft (20, 21). Transplanted SCID mice can be further complemented with cellular components of the human adaptive immune system as we have previously reported (19, 20) to investigate human-specific immune response against SARS-CoV-2 in the human gut.

Taken together we expect that SARS-CoV-2 administered intraluminally or systemically will readily infect cells of human gut xenografts and that this platform can serve as an experimental model for coronavirus infection and biology.

## MATERIALS AND METHODS

### SCID mouse human intestinal xeno-transplant model

C.B-17/IcrHsd-Prkdcscid (abbreviated as SCID) mice were purchased from Envigo Israel (Rehovot, Israel). All mice were housed in a pathogen-free facility, in individually ventilated cages (IVC), given autoclaved food and water. All animal use was in accordance with the guidelines and approval of the Animal Care and Use Committee (IACUC) of the Hebrew University of Jerusalem. IRB and IACUC approvals were obtained prospectively (Ethics Committee for Animal Experimentation, Hebrew University of Jerusalem; MD-11-12692-4 and the Helsinki Committee of the Hadassah University Hospital; 81-23/04/04).

Women undergoing legal terminations of pregnancy gave written, informed consent for use of fetal tissue in this study. Human fetal small bowel 12-18 weeks gestational age was implanted subcutaneously on the dorsum of the mouse as described previously (21). All surgical procedures were performed in an aseptic working environment in a laminar flow HEPA-filtered hood with isoflurane inhalation anesthesia (1.5 to 2% v/v isofluorane in O_2_). Before surgery, carprofen (5 mg/kg, Rimadyl, Pfizer Animal health) was administered subcutaneously. The surgical area was shaved and depilated (Nair hair removal cream) and the skin was scrubbed and disinfected with betadine and 70% (v/v) ethanol. After surgery the mice were provided with enrofloxacin-medicated water (Bayer Animal HealthCare AG) for 7 days and were closely monitored once a day for behavior, reactivity, appearance and defecation. Grafts developed in situ for 12-16 weeks prior to manipulation.

#### Histology and Immunofluorescence

As we have previously described (15, 20), human fetal gut and xenograft tissues were harvested for immunofluorescence staining and microscopic analysis. Tissues were fixed in 2.5% PFA overnight at room temperature, incubated with 15% (w/v) sucrose for 12 hours at 4°C and frozen in Tissue-Tek® (EMS, Hatfield, PA) embedding medium. Serial 10 μm cryosections were stained with phalloidin (abcam, ab176759) and 4’,6’-diamidino-2-phenylindole (DAPI) (Sigma, D9542). For immunofluorescence staining primary antibodies were anti-ACE2 (Invitrogen, #MA5-32307) and anti-TMPRSS2 (Santa Cruz, sc-515727). Alexa Fluor 594 conjugated goat anti-mouse IgG (Invitrogen, #A-11005) and Alexa Fluor 488 conjugated goat anti-rabbit IgG (Invitrogen, #A-11034) were used as secondary antibodies. Sections were mounted with VectaShield (Vector Laboratories, Burlingame, CA) and imaged with an Axio Imager M1 upright light microscope (Zeiss, Germany) coupled to a MR3 CCD camera system (ZEN 2012).

### Experimental model of systemic inflammatory response syndrome

Transplanted mice were subjected to systemic LPS (from *E. coli* serotype O55:B5, Sigma, Rehovot, Israel) at 0.1, 1 or 10 mg/kg by intraperitoneal injection. Viability and clinical signs were monitored and scored by visual examination (Fig. S5). Clinical score was adopted from Bougaki, et al (55). Four hours after LPS treatment mice were humanely sacrificed, and human gut transplants and mouse ileum were harvested. Full thickness sections and mucosal scrapings were collected for extraction of RNA.

### RNA sequencing analysis

RNA from LPS-treated (n ≥ 3) and PBS-treated (n ≥ 3) control human gut xenografts and mouse jejunum were purified using GenElute Mammalian Total RNA kit (Sigma) following manufacturer’s instructions, and RNA samples were quantified using a NanoDrop ND-1000 spectrophotometer (Thermo Fisher Scientific Inc., Wilmington, DE, US). A library was made with the TruSeq Stranded total RNA library prep kit according to the manufacturer’s description (Illumina inc, San Diego, USA) and RNA was sequenced on Illumina HiSeq2500 using Illumina stranded TruSeq protocol. An average of ~30 million 125-nt paired-end reads was sequenced for each sample.

The read quality of all samples were assessed separately using FASTQC (http://www.bioinformatics.bbsrc.ac.uk/projects/fastqc) and raw reads were pre-processed using Cutadapt (19) to remove adapter and low quality sequences. The paired end reads were aligned using TopHat 2.0.14 to a reference genome that consisted of the combination of the reference genomes GRCh38 (human, Ensembl release 83) and GRCm38 (mouse, Ensembl release 83), modifying the chromosome names in order to identify the two species. The alignments for the two species were then separated with the original chromosome names restored. HTSeq-count 0.6.1p1 was then used to produce read counts for each gene based on the reference annotations Homo_sapiens.GRCh38.80.gtf and genes_mm10_ens.gtf.

Adjusted log2 fold change and adjusted p-values (corrected for multiple tests) were calculated using DESeq2 (56) to determine significant differences in gene expression. Genes were scored according to an absolute fold-change ≥ 1 and adjusted p-value of < 0.05.

### Ortholog selection

The Ensembl Biomart (57) was used to extract all orthologous mouse genes for all genes in the human genome GRCh38 (release 83). The result was a list of orthologs with a one-to-one, one-to-many or many-to-many relationship with the human gene, and also a binary confidence value for each ortholog. To select the best match ortholog for each human gene the following procedure was followed. All one-to-one orthologs were selected and retained. For one-to-many and many-to-many orthologs, where there was only one high confidence ortholog (confidence = 1) this was selected. If there was greater than one or no high confidence orthologs the ortholog with the highest percentage identity with the mouse gene was selected, else if the percentage identity was the same for each ortholog the ortholog with the matching gene name was selected and retained. This combined list resulting from this procedure consisted of 18,719 murine orthologs. To keep the text concise and readable we use uppercase HGNC gene names throughout when referring to orthologs in both human and mouse.

### Gene Set Enrichment Analysis

GSEA was performed using the using the fgsea package (version 1.2.2) in R (29). The ImmuneSigDB Gene Sets (c7.all.v6.1.symbols.gmt) were downloaded from the Broad Institute website (http://software.broadinstitute.org/gsea/msigdb/collections.jsp). Gene Ontology analysis to compare a target set of genes of interest with the background set of genes from mouse or human (GRCh38 or GRCm38) used GOrilla tool (58).

## ACKNOWLEDGEMENTS

This work was supported by funding from the European Union Seventh Framework Programme (FP7/2012-2017) under grant agreement no. 305564 as partners of the SysmedIBD research consortium (to Werner Muller, University of Manchester, United Kingdom).

The technical assistance of Dr. Irit Shoval in the preparation of gut transplants and LPS challenge studies is acknowledged.

## AUTHORS’ CONTRIBUTIONS

NYS designed and supervised the study. LM, FB, ND, HM, RST, PR, DR, VAPMS and NYS contributed to the genome assembly, annotation and analyzed the transcriptomic data. ENE, EN, NM, IS and SY, contributed to the experiments and their setup. NYS and LM wrote the first versions of the manuscript with contribution from all authors. All authors assisted with editing of the manuscript and approved the final version.

---

**Fig. S1.**
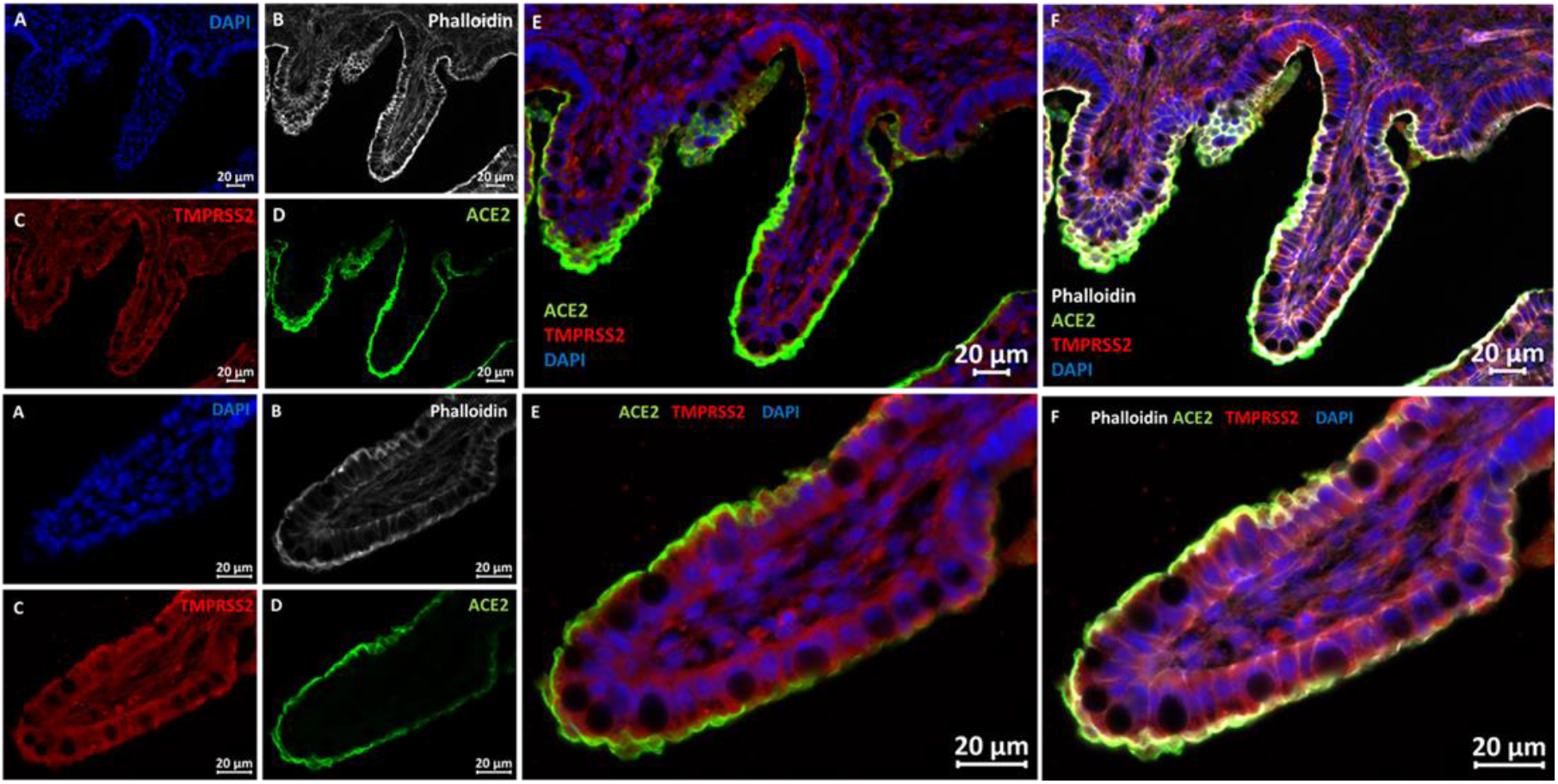
SARS-CoV-2 entry molecule; ACE2 and TMPRSS2 are co-expressed in the mucosal villi of the gut of 17 weeks old human fetus. Fluorescence microscopy was used to analyze cryosections of normal human fetal gut stained using DAPI, phalloidin and anti ACE2 and TMPRSS2 antibodies. Microvilli of gut epithelial cells are demonstrated by phalloidin staining co-localized with ACE2 staining. Scale bars 20 μm.

**Fig. S2.**
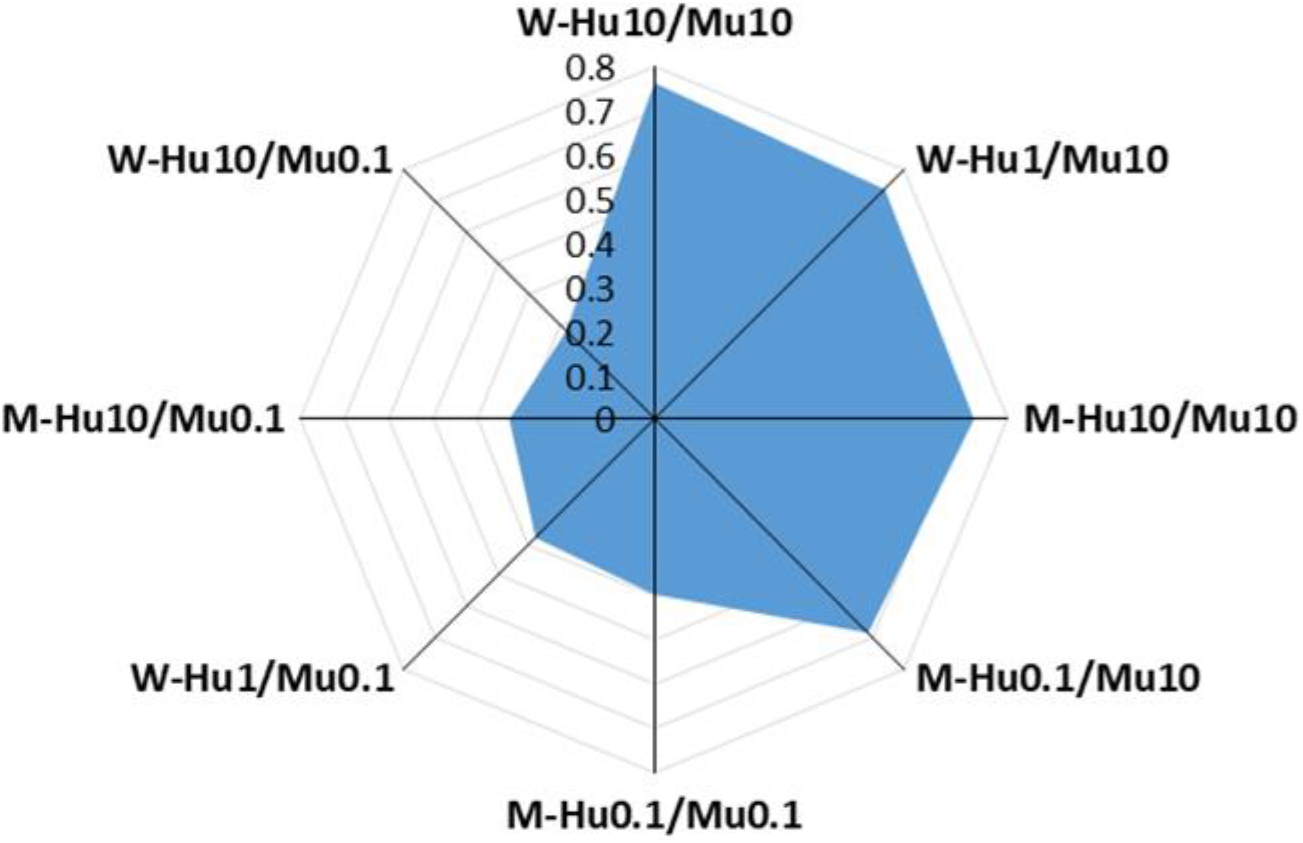
Genomic response in the mouse gut induced by high dose LPS correlate best with genomic response in the human gut. Radar plot of Spearman correlations (ρ) described in Fig. 4. Diagram shows correlations between genomic response in whole (W) or mucosal (M) samples obtained from human (Hu) or murine (Mu) gut treated with 0.1, 1 or 10 mg/kg LPS.

**Fig. S3.**
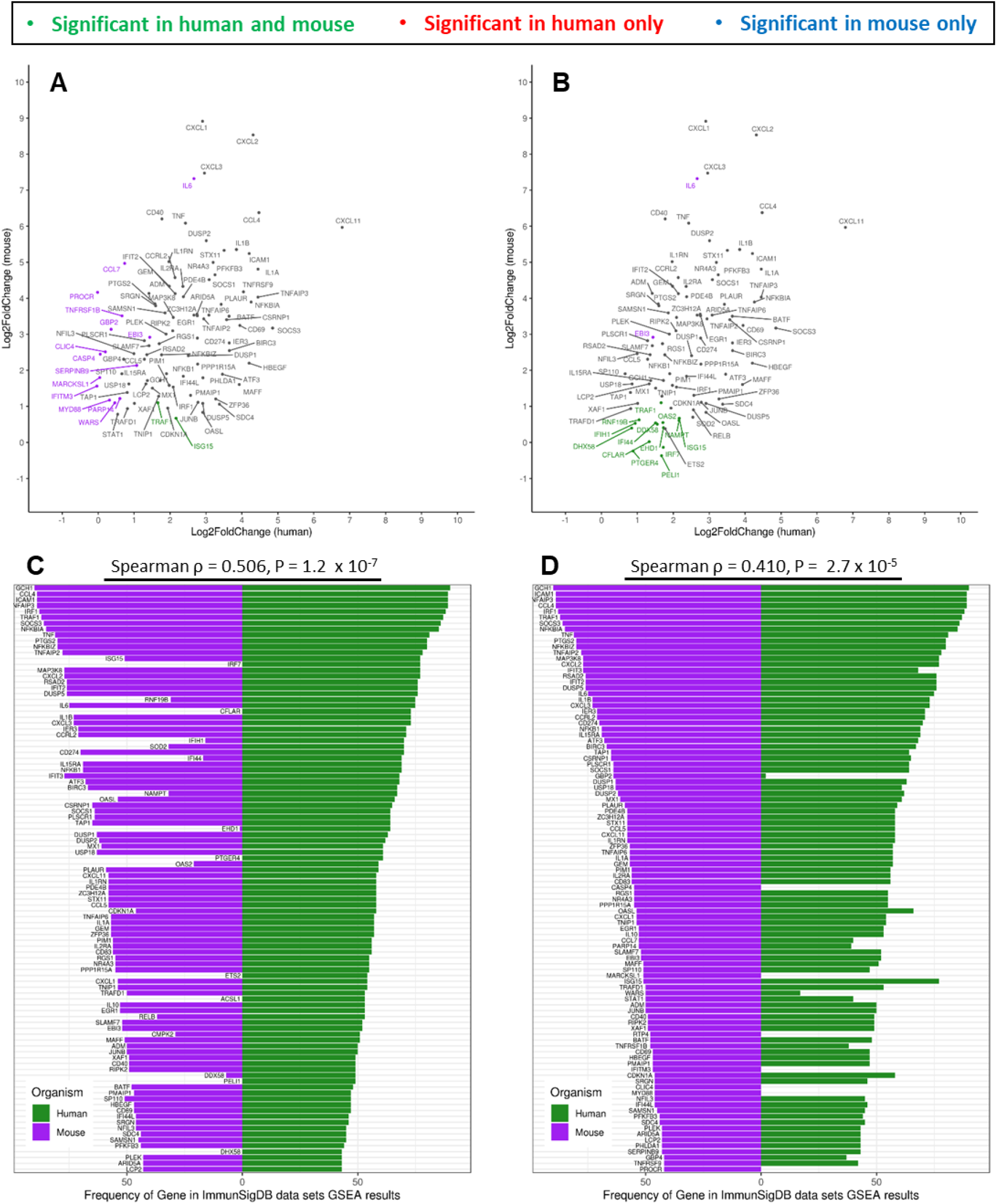
(A) Comparison of log fold changes in the top 100 leading edge genes from the most enriched genes sets in mouse, compared to human (B) Comparison of log fold changes in the top 100 leading edge genes from the most enriched genes sets in human, compared to mouse (C) Frequency histogram of the top 100 genes occurring in the leading edge of the gene sets enriched in mouse, compared to human. (D) Frequency histogram of the top 100 genes occurring in the leading edge of the gene sets enriched in mouse, compared to human.

**Fig. S4.**
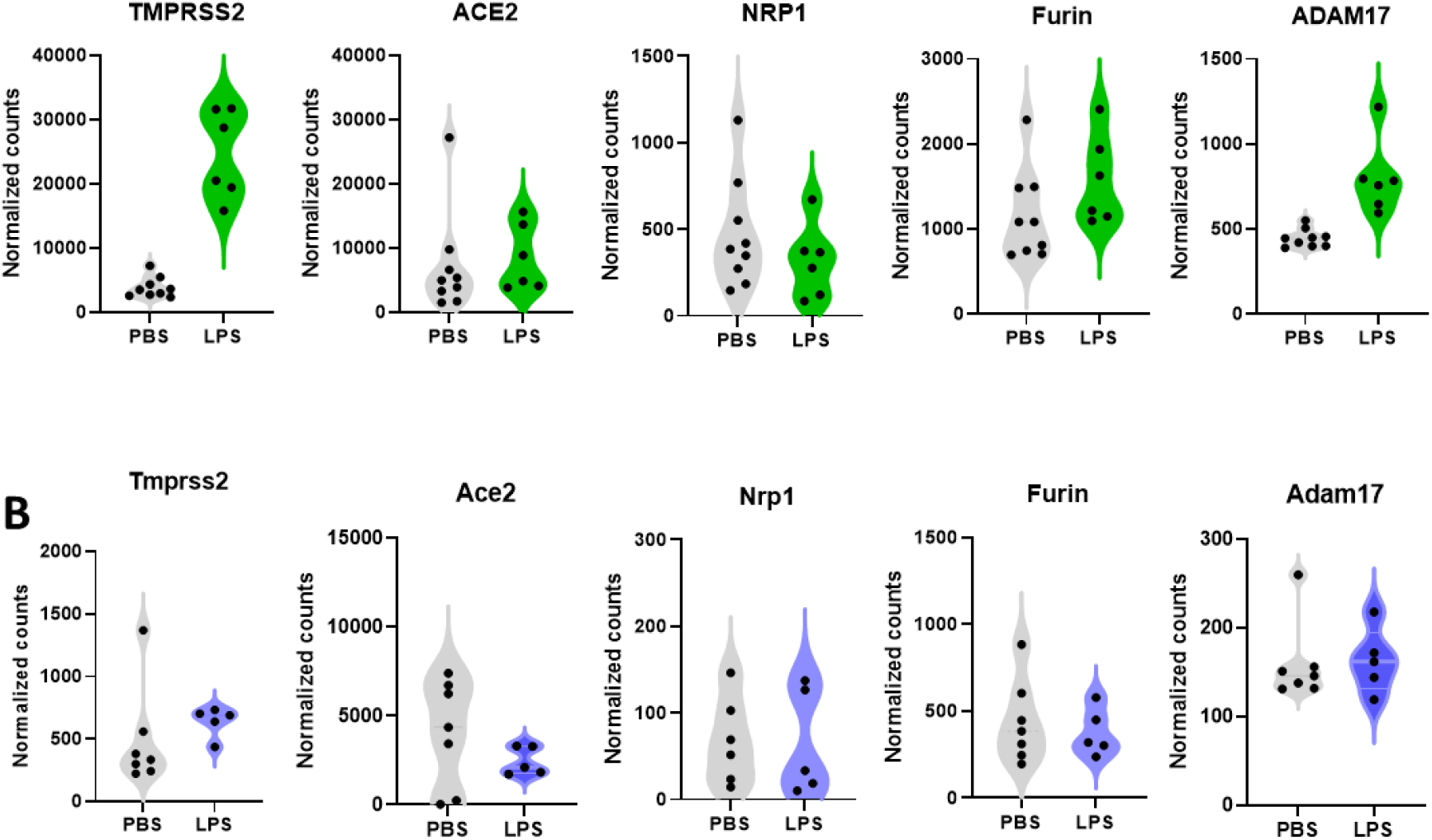
TMPRSS2 and ADAM17 are upregulated in LPS-treated human gut xenografts but not in host mouse gut. Violin plots showing normalized and scaled expression of TMPRSS2, ACE2, NRP1, Furin and ADAM17 in human gut xenografts (A) and host mouse gut (B) from PBS- and LPS-treated (10 mg/kg) mice.

**Fig. S5.**
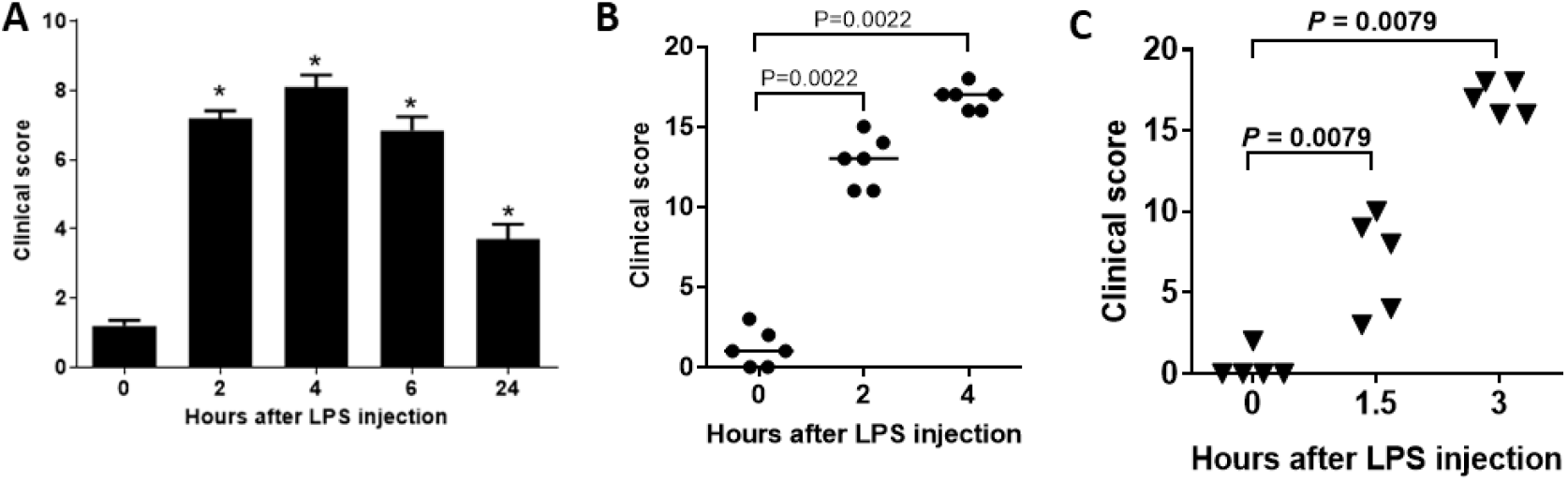
Clinical score of SCID mice with mature human gut xenografts treated by a single intraperitoneal injection of 0.1 (A), 1 (B) and 10 (C) mg/kg LPS. Mice were evaluated clinically and scored before treatment and at the indicated time points after LPS treatment (A-C). Parametric and non-parametric data were calculated as the mean ± s.d. (A) and median (horizontal bars in B-C) respectively. For comparison of parametric and non-parametric data, t- test and non-parametric Mann–Whitney two-independent-samples test (respectively) were applied using GraphPad Prism 6 (GraphPad Software, Inc.). A P value of 0.05 or less was considered significant. A; mean score (*, P < 0.05) significantly different compared to time point 0.

## Notes

### Competing Interest Statement

The authors have declared no competing interest.

